# Cognitive Control in Pediatric Cancer Survivors: A task-fMRI study

**DOI:** 10.64898/2026.06.23.734009

**Authors:** Siddharth Nayak, Shounak Nandi, Faye McKenna, Sonya Henry, Tim Q. Duong

## Abstract

**Background:** Chemotherapy-related cognitive impairment is a well-documented concern among cancer survivors, yet the neural mechanisms underlying deficits in cognitive control remain poorly understood. This study examined group differences in brain activation during a flanker task using functional MRI (fMRI) between chemotherapy-exposed participants and healthy controls.

**Methods:** Participants (21 survivors (24.9 years old; 71.4 % female; 15 years from diagnosis) and 21 healthy controls (26.7 years old; 61.9 % female) completed a flanker task during fMRI, with congruent and incongruent conditions. Reaction time, accuracy, and Flanker scores were collected. Whole-brain group comparisons were performed for congruent, incongruent, and incongruent > congruent contrasts. Associations between the incongruent > congruent contrast and cognitive performance were examined.

**Results:** Compared to controls, the Chemo group had longer reaction times in both congruent and incongruent conditions (p < .001) and lower NIH Flanker scores (p = .01), with no differences in accuracy. They showed reduced activation in the bilateral inferior frontal gyri, supplementary motor area, and bilateral caudate, but greater activation in the right inferior temporal and cerebellar regions. The incongruent > congruent contrast correlated with increased activation in the orbitofrontal cortex, inferior temporal gyri, and fusiform gyrus with cognitive performance.

**Conclusions:** Chemotherapy-exposed participants showed cognitive control deficits and altered neural activation during a flanker task, indicating disrupted recruitment of frontoparietal and subcortical regions key for conflict processing. These findings improve understanding of neural causes of chemotherapy-related cognitive impairment and may help identify at-risk survivors and guide personalized rehabilitation.

## 1 Introduction

Treatments of pediatric acute lymphoblastic leukemia (ALL), the most common childhood malignancy, achieve 5-year survival rates exceeding 90% (Malard and Mohty 2020; Hunger and Mullighan 2015). This remarkable therapeutic success has been achieved through intensive chemotherapy regimens, which include high-dose intravenous and intrathecal methotrexate and other agents (Malard and Mohty 2020; Hunger and Mullighan 2015). However, the growing population of long-term survivors faces significant neurocognitive sequelae that adversely impact educational attainment, vocational success, and quality of life (Jacola et al. 2025; Cheung and Krull 2015; Chendan Zhou et al. 2020b).

Chemotherapy induces oxidative stress, neuroinflammation, mitochondrial dysfunction, and white matter damage, which could collectively compromise the structural and functional integrity (Park et al. 2025; Chughtai et al. 2025), which could result in chemotherapy-related cognitive impairment (CRCI) (Chughtai et al. 2025) spanning multiple neurocognitive domains, with attention, executive function, processing speed, working memory, and intelligence being particularly vulnerable (Jacola et al. 2025; Cheung and Krull 2015; Chendan Zhou et al. 2020b). Current clinical practice guidelines from the National Comprehensive Cancer Network (NCCN) recommend neurocognitive monitoring during and after completion of therapy for all pediatric cancer patients, given the established risk for neurocognitive late effects associated with CNS-directed chemotherapy (Inaba et al. 2025; Gajjar et al. 2025). Monitoring should occur at treatment completion and/or at school entry or re-entry, with baseline assessment considered to provide context for appreciating change (Inaba et al. 2025).

Many neuroimaging studies have shown cortical thinning, reduced gray and white matter volumes, altered white-matter microstructure in ALL patients as well as resting state functional connectivity compared to controls (see review papers (Gandy et al. 2021; C. Zhou et al. 2020a)). In contrast, only a few task-based functional magnetic resonance imaging (fMRI) studies have been reported to date in this population. Robinson et al. used N-back task and showed greater activation in dorsolateral/ventrolateral prefrontal cortex and anterior cingulate during working memory tasks (Robinson et al. 2010). Fallah et al. used attention task and showed increased activation in ventral frontal, insula, and caudate regions during alerting tasks (Fellah et al. 2019). This compensatory recruitment suggests that the developing brain attempts to overcome neural inefficiency by engaging additional cognitive resources. Stefancin et al. used N-back task and examined neural correlates during both successful and unsuccessful task performance (Stefancin et al. 2020) and found that survivors exhibited reduced posterior cingulate activation during correct responses but greater angular gyrus activation during errors and greater superior parietal activation during no-response trials, suggesting that working memory impairment stems specifically from an inability to manipulate and retrieve information from memory rather than a global processing deficit.

Other task-fMRI studies evaluated factors that influence task-evoked activations. Edelmann et al. showed dexamethasone exposure is associated with greater memory impairment and altered hippocampal-related activation compared with prednisone using a word-recognition task and (Edelmann et al. 2013). Gandy et al. found no statistically significant associations between genetic variants and task-fMRI BOLD activity (Gandy et al. 2023) but working memory network by fMRI was more impaired in female survivors than male survivors (Gandy et al. 2022). Conklin et al. found significant pre- to post-training reduction in activation of left lateral prefrontal and bilateral medial frontal areas using a spatial working-memory task (Conklin et al. 2015). In an episodic encoding task, whole-brain statistical map analysis revealed increased blood oxygenation level-dependent signal/activation in several brain regions during unsuccessful encoding in ALL survivors, potentially reflecting ineffective neural recruitment (Monje et al. 2013). These task-based fMRI studies employed a heterogeneous array of cognitive paradigms and fMRI designs their findings are heterogeneous.

The Flanker task is a validated experimental paradigm for assessing cognitive control and executive function in healthy adults (McDermott et al. 2017). This task measures the ability to selectively attend to target stimuli while inhibiting responses to flanking distractors, thereby engaging neural networks critical for conflict detection, response inhibition, and attentional control. Performance on cognitive neuroscience paradigms like the Flanker task provides complementary information to standardized neuropsychological measures, potentially offering more sensitivity to subtle executive function deficits and mechanistic insights into the neural substrates of CRCI (Hooke et al. 2021).

Building on prior studies, we performed fMRI while subjects performed the Flanker task in the MRI scanner to further investigate CRCI in a diverse population of pediatric cancer patients. Additional neurocognitive tests performed outside the scanners and correlation analysis was performed with the fMRI activations. By examining both neural activation patterns and behavioral performance across congruent and incongruent trial conditions, this investigation aims to elucidate the specific alterations in frontoparietal attention networks, default mode network dynamics, and compensatory neural mechanisms that characterize CRCI in this vulnerable population. Understanding the neural substrates is essential for developing targeted interventions, identifying high-risk patients through early biomarkers, and ultimately improving long-term neurocognitive outcomes in the growing population of pediatric hematologic cancer survivors.

## 2 Methods

### 2.1 Participant Recruitment

The study was approved by the Institutional Review Board of Albert Einstein College of Medicine (IRB-2020-12265), and all participants and/or their legal guardians provided written informed consent. Ethical principles were strictly adhered to, including maintaining participant anonymity and minimizing potential risks associated with the research.

Participants (aged 15-35 years) included pediatric hematologic cancer survivors (chemo group) and age-matched healthy controls (control group). Cancer patients were childhood cancer survivors who were 10-25 years post-chemotherapy. Eligible cancer types included acute lymphoblastic leukemia (ALL), acute myeloid leukemia, neuroblastoma, non-CNS solid tumors, and Hodgkin lymphoma.

### 2.2 Magnetic Resonance Imaging

MRI scans were acquired in a 3 Tesla Philips scanner equipped with a 32-channel head coil (Elition 3.0 T X, Philips Medical Systems, Best, The Netherlands) at the Gruss Magnetic Resonance Research Center of Albert Einstein College of Medicine, New York. A high-resolution T1-weighted whole head structural image was acquired using axial 3D-MPRAGE parameters over a 240 mm FOV and 1.0 mm isotropic resolution, TE=4.6 ms, TR=9.9 ms, TI=806 ms. The blood oxygen level-dependent (BOLD) images for task-based functional MRI (fMRI) were acquired using single-shot echo-planar imaging, with a 224 x 224 mm field of view (FOV) and 2 mm isotropic resolution; parameters were TE = 28 ms, TR = 2000 ms, flip angle = 90 °, and 37 trans-axial slices. Functional MRI data for the Flanker task (Zhu, Zacks, and Slade 2010) were acquired in an event-related design in a single run, approximately 5 minutes long and comprising 144 volumes distributed across congruent, incongruent, and neutral conditions. The stimulus was a row of five arrows (e.g., <<<<< or >><>>), and participants had to respond by pressing the button to the middle arrow as the target. The arrows on either side of the middle arrow are distractors (flankers). In the congruent condition, flankers point in the same way as the target (e.g., <<<<< or, >>>>>), making it easier to respond. In the incongruent condition, flankers point in the opposite way as the target (e.g., <<><< or, >><>>), creating interference, hence making it difficult to respond. Lastly, there was a neutral condition where the flankers were of irrelevant shapes (e.g., -->-- or, ++<++), added as null trials of no interest. Participants were instructed to press the button on a response pad corresponding to the direction of the central target arrow on all trials; reaction time (RT) and accuracy (ACC) were recorded. Although our focus has been on congruent, incongruent, and incongruent > congruent contrasts, future research would also examine neutral conditions, both independently and in comparison to these other contrasts.

### 2.3 fMRI Processing

Functional MRI images were preprocessed using FSL (Version 6.0.5) (Jenkinson et al. 2012), FMRIB’s Software Library (http://fsl.fmrib.ox.ac.uk/fsl), and consisted of non-brain removal using BET, motion correction with MCFLIRT, slice-timing correction for interleaved acquisitions using Fourier-space time-series phase shifting, geometric distortion correction using B0 images, high pass temporal filtering using Gaussian-weighted least-squares straight line fitting (r=100 s), spatial smoothing using a Gaussian kernel with full-width half-maximum 5 mm, affine co-registration to high-resolution T1-weighted images, and normalization to standard space (Montreal Neurological Institute atlas, 2 mm template) using a combination of affine and nonlinear registration. The first-level analysis included modeling condition-specific task timing files using a generalized linear model (GLM) in FSL FEAT to account for task effects, and estimating motion artifacts using MCFLIRT (Smith et al. 2004). We created contrasts for the congruent, incongruent, and incongruent > congruent conditions, while treating neutral trials as the baseline. Due to the very fast event-related design, events were modeled as delta (stick) functions at stimulus onset and convolved with a canonical double-gamma hemodynamic response function (HRF) to generate regressors for the GLM. Higher-level analyses were performed to generate contrast maps for congruent, incongruent, and incongruent > congruent contrasts using mixed-effects (Flame 1+2).

### 2.4 NIH Toolbox: Cognition Battery

Cognitive performance was assessed using the NIH Toolbox Cognition Battery (Weintraub et al. 2013), which included the Fluid Intelligence and Early Childhood composite scores, the Flanker Inhibitory Control and Attention test, and the Pattern Comparison Processing Speed tests. We included the Flanker Inhibitory Control (Shono et al. 2024) test in the current study to compare with the in-scanner flanker task.

### 2.5 Statistical Analyses

Descriptive statistics were used to summarize all study measures (mean ± SD for continuous variables; counts and percentages for categorical variables). Voxel-wise analyses were conducted on the fMRI contrast maps in MNI space using FSL randomize with 5000 permutations and a p < .01, whole-brain-corrected, cluster-forming family-wise error (FWE)-corrected threshold. Group-level comparisons were conducted across groups (Chemo, Control, and Control vs. Chemo) to assess differences in activation/deactivation patterns for congruent, incongruent, and incongruent > congruent contrasts. The group-level maps were associated with within-scanner (RT) and outside-scanner (NIH Flanker task) behavioral variables, after adjusting for covariates (age and sex) for the incongruent > congruent contrast. Additionally, first-level activations and deactivations within the dorsal attention network (DAN) and the default mode network (DMN) were computed, and box plots were generated using customized Python scripts to compare the Chemo and Control groups across congruent, incongruent, and incongruent > congruent contrasts. Statistical analyses were performed in SPSS (version 29; SPSS, Inc., Chicago, IL), and tests were two-sided; *p < 0.05* was considered statistically significant. Functionally distinct clusters for the target contrasts after whole-brain cluster correction were created using the AAL3 atlas (Rolls et al. 2020). Brain maps were visualized using BrainNet Viewer (www.nitrc.org/projects/bnv/).

## 3 Results

### 3.1 Subject profiles and behavioral results

Demographic characteristics for the two groups are described in **Table 1**. There were no significant group differences in age and sex; however, groups differed significantly in race (p = .0025) and ethnicity (p = .0026), with the Chemo group having a higher proportion of African American and Hispanic participants. Behavioral data (RT and ACC) were unavailable for 3 controls in the congruent condition and 1 control in the incongruent condition. However, all participants were retained for the imaging analyses described below. The Chemo group had significantly longer RT than the Control group in both the congruent (p < .001) and incongruent (p < .001) conditions, but there was no group difference in accuracy between conditions. The Chemo group had significantly lower scores on the NIH Flanker task compared to the Control group (p = .01).

**Table 1:**
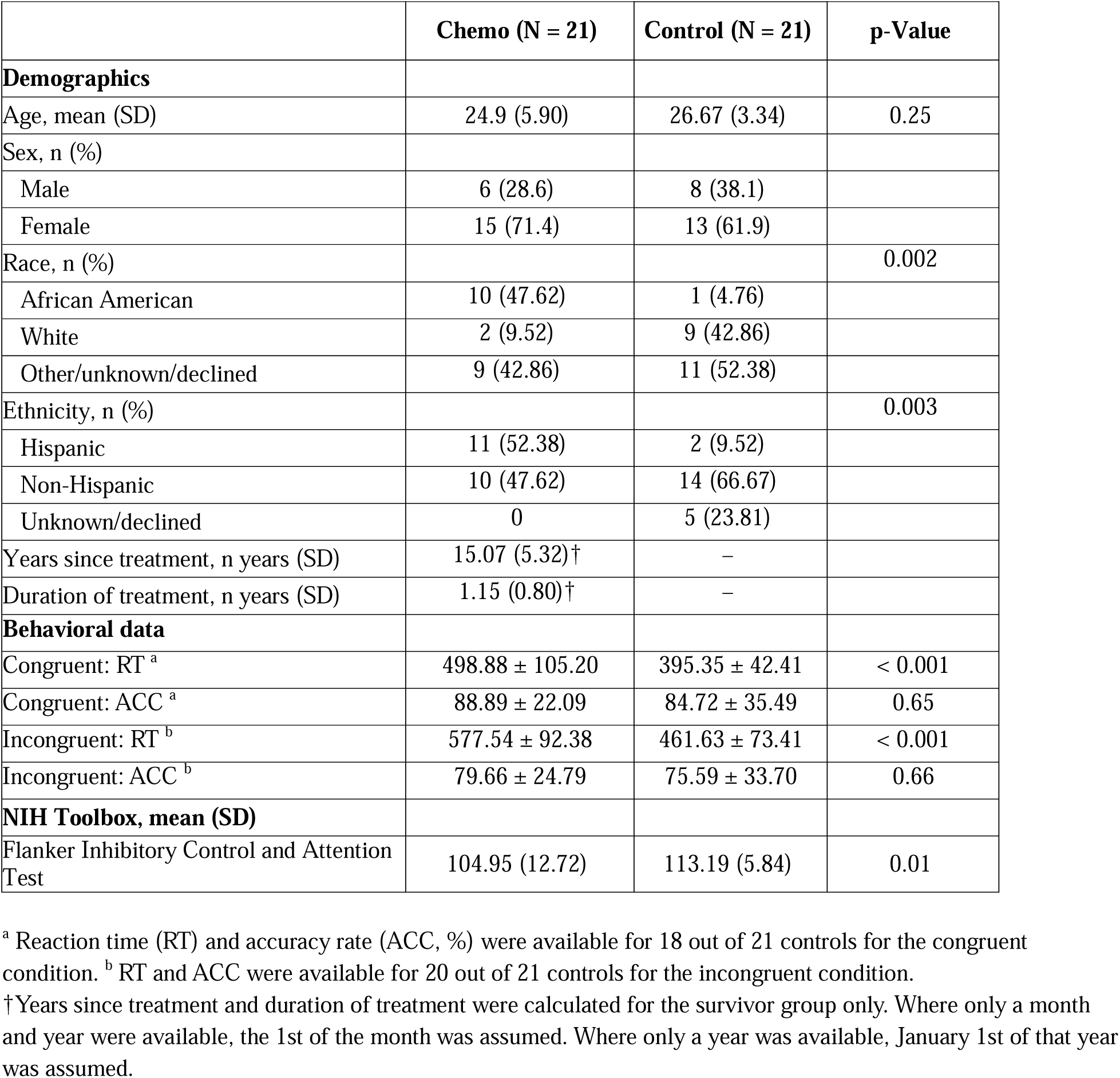
Sample characteristics.

### 3.2 Group differences in Congruent, Incongruent, and Incongruent > Congruent contrasts

**Table 2A** and **Figure 1A** present the results for the congruent contrast. Compared with the Chemo group, the Control group showed greater activation in the left hemisphere, specifically in the middle/inferior temporal gyri (MTG/ITG), supplementary motor area (SMA), and superior frontal gyrus (SFG), whereas lesser activation in the right MTG and right inferior frontal gyrus (IFG). **Table 2B** and **Figure 1B** present the results for the incongruent contrast. Compared with the Chemo group, the Control group showed greater activation in bilateral IFG, left MTG, and right insula, whereas lesser activation in the right ITG and right IFG/MTG. **Table 2C** and **Figure 1C** present the results for the incongruent > congruent contrast. Compared with the Chemo group, the Control group showed greater activation in the bilateral caudate, right superior temporal gyrus (STG)/MTG, and right postcentral gyrus for the incongruent > congruent contrast, whereas lesser activation in the left ITG and bilateral cerebellum. The complete set of activations and deactivations for the congruent, incongruent, and incongruent > congruent contrast is presented in **Table S1**.

**Figure 1:**
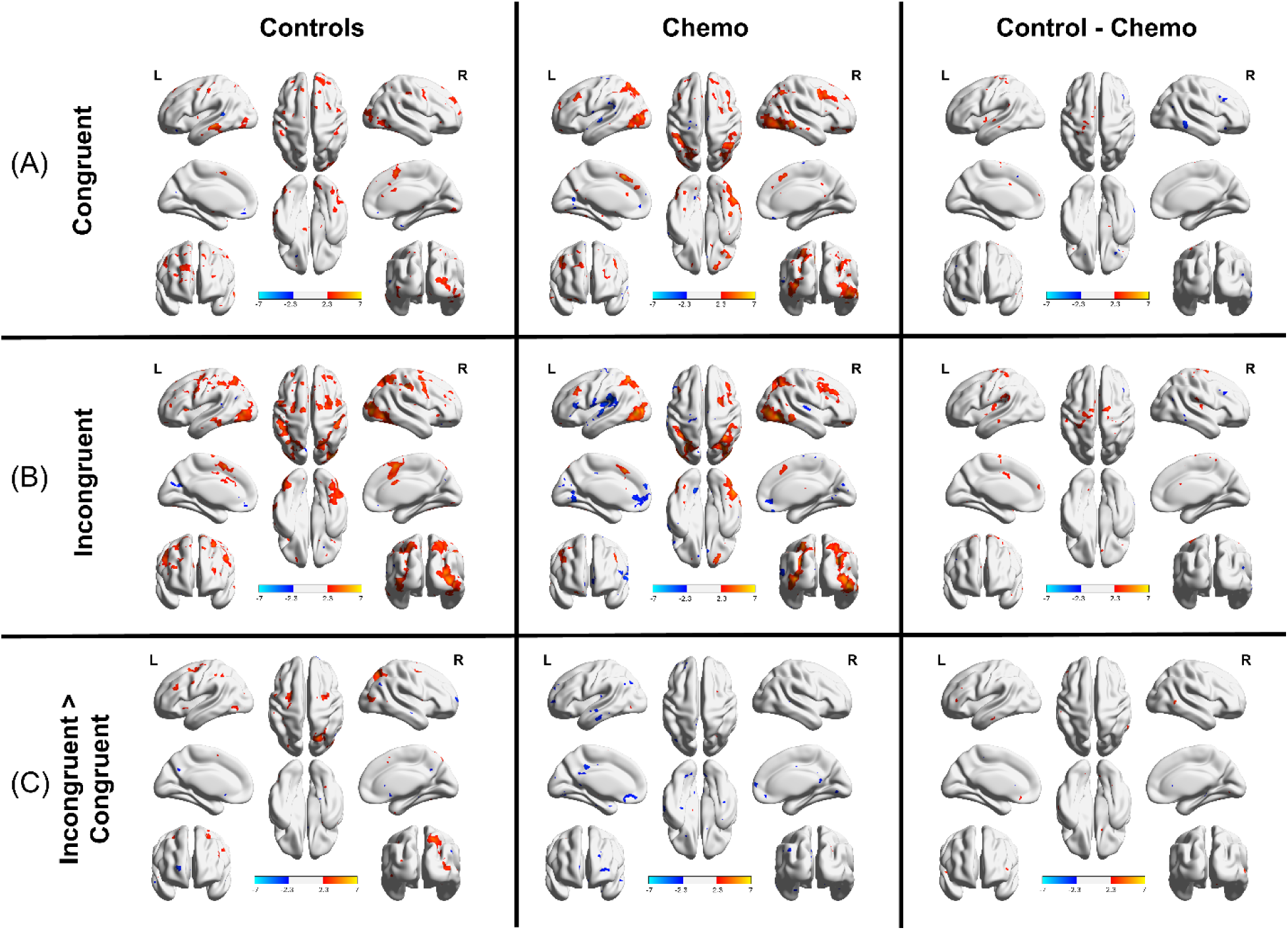
Group-level analysis of Controls, Chemo, and Controls – Chemo group for (A) Congruent, (B) Incongruent, and (C) Incongruent > Congruent contrasts. Activation (red) and deactivation (blue) regions across the whole brain. Significant results (p < .01 FWE-corrected) are shown on a MNI152 surface template.

**Table 2:**
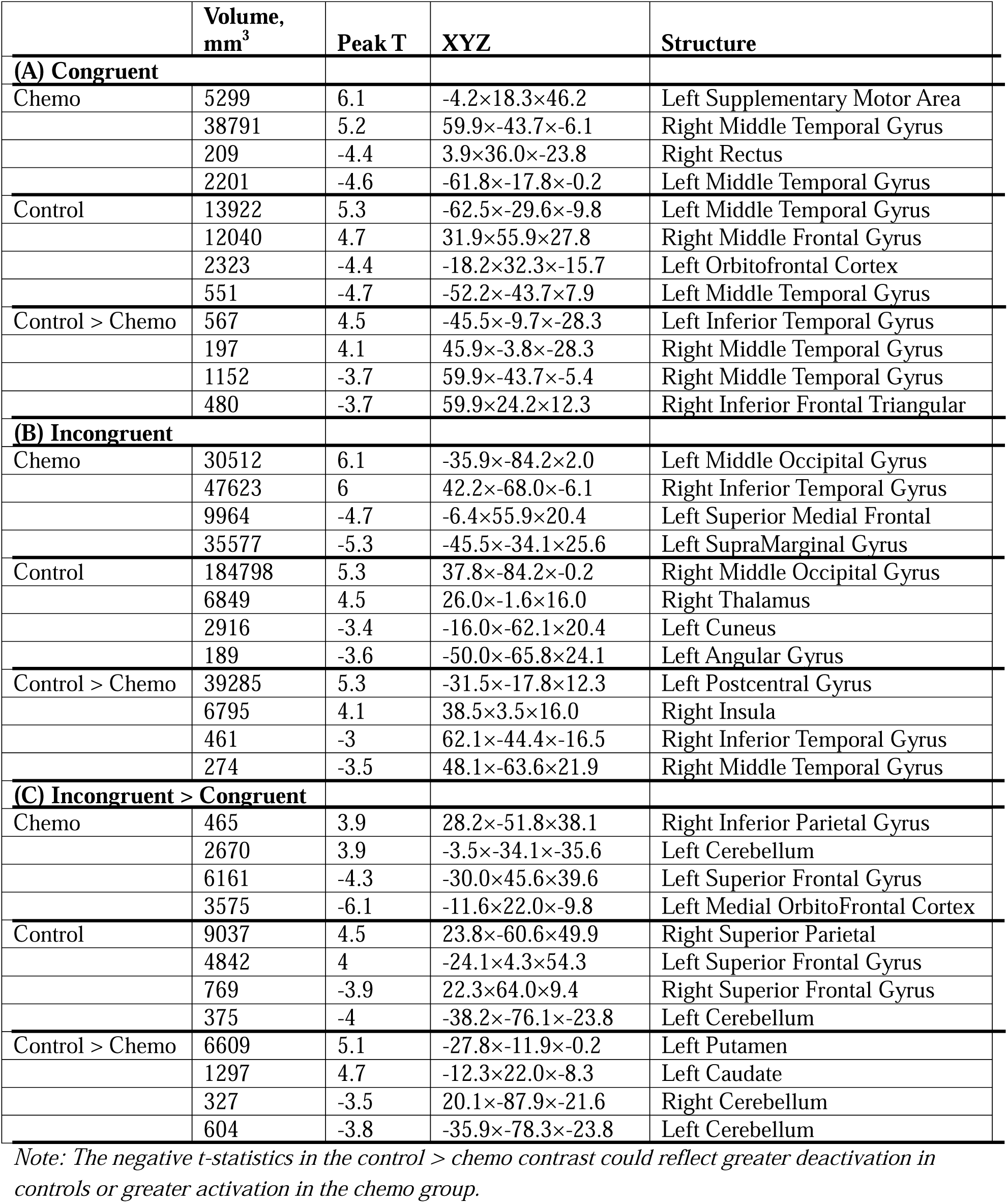
Brain regions showing significant activation/deactivation for the (A) Congruent, (B) Incongruent, and (C) Incongruent > Congruent contrasts between Chemo and Control groups.

|  | Volume,<br>mm<sup>3</sup> | Peak T | XYZ | Structure |
| --- | --- | --- | --- | --- |
| **(A) Congruent** |  |  |  |  |
| Chemo | 5299 | 6.1 | -4.2×18.3×46.2 | Left Supplementary Motor Area |
|  | 38791 | 5.2 | 59.9×-43.7×-6.1 | Right Middle Temporal Gyrus |
|  | 209 | -4.4 | 3.9×36.0×-23.8 | Right Rectus |
|  | 2201 | -4.6 | -61.8×-17.8×-0.2 | Left Middle Temporal Gyrus |
| Control | 13922 | 5.3 | -62.5×-29.6×-9.8 | Left Middle Temporal Gyrus |
|  | 12040 | 4.7 | 31.9×55.9×27.8 | Right Middle Frontal Gyrus |
|  | 2323 | -4.4 | -18.2×32.3×-15.7 | Left Orbitofrontal Cortex |
|  | 551 | -4.7 | -52.2×-43.7×7.9 | Left Middle Temporal Gyrus |
| Control > Chemo | 567 | 4.5 | -45.5×-9.7×-28.3 | Left Inferior Temporal Gyrus |
|  | 197 | 4.1 | 45.9×-3.8×-28.3 | Right Middle Temporal Gyrus |
|  | 1152 | -3.7 | 59.9×-43.7×-5.4 | Right Middle Temporal Gyrus |
|  | 480 | -3.7 | 59.9×24.2×12.3 | Right Inferior Frontal Triangular |
| **(B) Incongruent** |  |  |  |  |
| Chemo | 30512 | 6.1 | -35.9×-84.2×2.0 | Left Middle Occipital Gyrus |
|  | 47623 | 6 | 42.2×-68.0×-6.1 | Right Inferior Temporal Gyrus |
|  | 9964 | -4.7 | -6.4×55.9×20.4 | Left Superior Medial Frontal |
|  | 35577 | -5.3 | -45.5×-34.1×25.6 | Left SupraMarginal Gyrus |
| Control | 184798 | 5.3 | 37.8×-84.2×-0.2 | Right Middle Occipital Gyrus |
|  | 6849 | 4.5 | 26.0×-1.6×16.0 | Right Thalamus |
|  | 2916 | -3.4 | -16.0×-62.1×20.4 | Left Cuneus |
|  | 189 | -3.6 | -50.0×-65.8×24.1 | Left Angular Gyrus |
| Control > Chemo | 39285 | 5.3 | -31.5×-17.8×12.3 | Left Postcentral Gyrus |
|  | 6795 | 4.1 | 38.5×3.5×16.0 | Right Insula |
|  | 461 | -3 | 62.1×-44.4×-16.5 | Right Inferior Temporal Gyrus |
|  | 274 | -3.5 | 48.1×-63.6×21.9 | Right Middle Temporal Gyrus |
| **(C) Incongruent > Congruent** |  |  |  |  |
| Chemo | 465 | 3.9 | 28.2×-51.8×38.1 | Right Inferior Parietal Gyrus |
|  | 2670 | 3.9 | -3.5×-34.1×-35.6 | Left Cerebellum |
|  | 6161 | -4.3 | -30.0×45.6×39.6 | Left Superior Frontal Gyrus |
|  | 3575 | -6.1 | -11.6×22.0×-9.8 | Left Medial OrbitoFrontal Cortex |
| Control | 9037 | 4.5 | 23.8×-60.6×49.9 | Right Superior Parietal |
|  | 4842 | 4 | -24.1×4.3×54.3 | Left Superior Frontal Gyrus |
|  | 769 | -3.9 | 22.3×64.0×9.4 | Right Superior Frontal Gyrus |
|  | 375 | -4 | -38.2×-76.1×-23.8 | Left Cerebellum |
| Control > Chemo | 6609 | 5.1 | -27.8×-11.9×-0.2 | Left Putamen |
|  | 1297 | 4.7 | -12.3×22.0×-8.3 | Left Caudate |
|  | 327 | -3.5 | 20.1×-87.9×-21.6 | Right Cerebellum |
|  | 604 | -3.8 | -35.9×-78.3×-23.8 | Left Cerebellum |

*Note: The negative t-statistics in the control > chemo contrast could reflect greater deactivation in controls or greater activation in the chemo group.*

### 3.3 Association of Incongruent > Congruent contrast with cognitive tests

**Table 3A** and **Figure 2A** present the association for the Incongruent > Congruent contrast with the reaction time derived from the fMRI flanker task. The Chemo group showed greater activation in the right OFC, left cerebellum, and right ITG, as well as greater deactivations in the right STG and left middle frontal gyrus (MFG). **Table 3B** and **Figure 2B** present the association for Incongruent > Congruent contrast with cognitive tests for the Flanker task. The Chemo group showed greater activation in the bilateral ITG, right OFC, and bilateral fusiform gyrus, as well as greater deactivations in the left temporal pole and left pallidum. The complete set of activations and deactivations for the incongruent > congruent contrast in association with cognitive tests is presented in **Table S2**.

**Figure 2:**
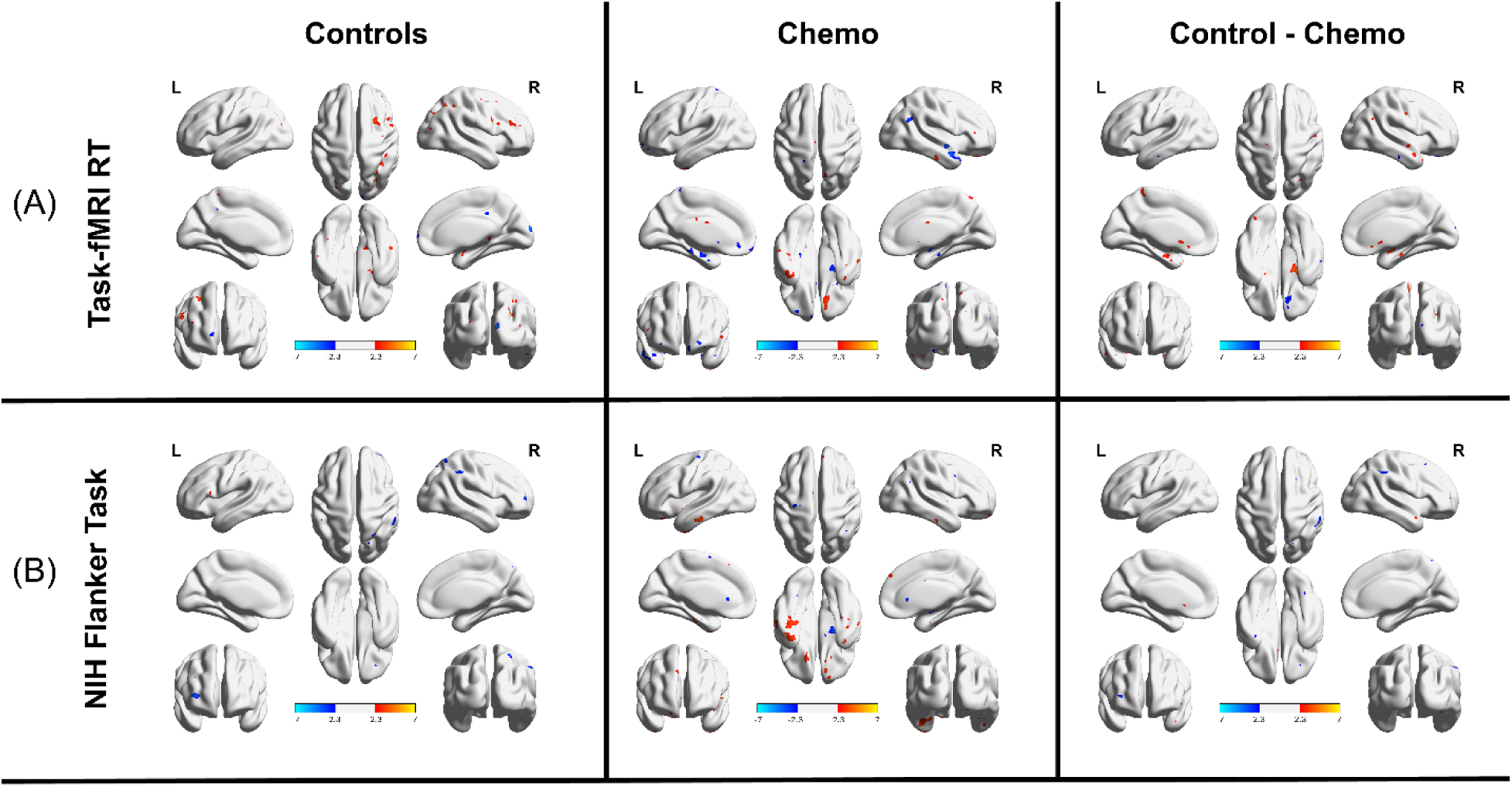
Association of Incongruent > Congruent contrast with (A) reaction time (RT) and (B) NIH flanker task scores across groups. Activation (red) and deactivation (blue) regions across the whole brain. Significant results (p < .01 FWE-corrected) are shown on a MNI152 surface template.

**Table 3:**
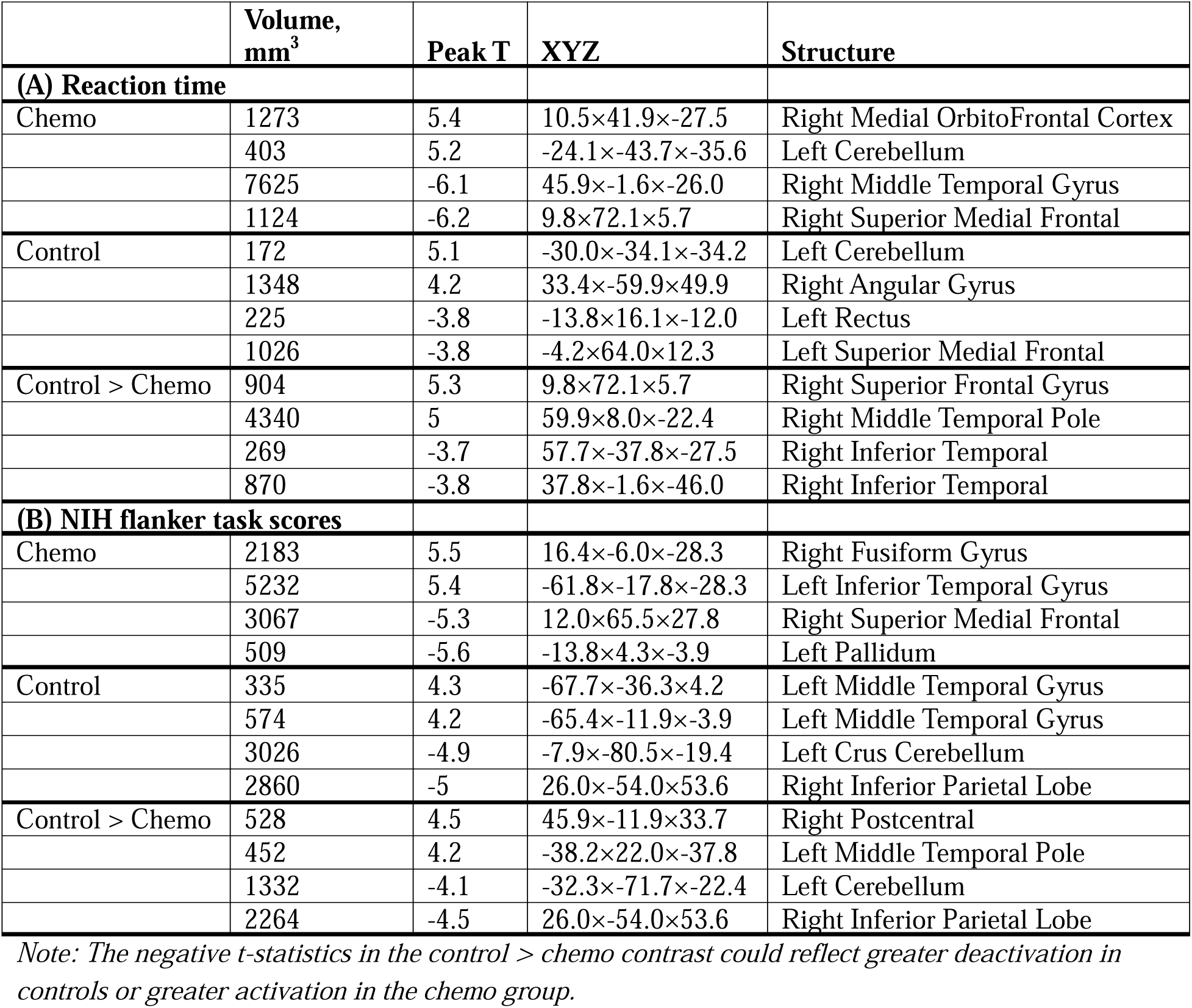
Brain regions showing significant association of Incongruent > Congruent contrast with (A) reaction time and (B) NIH flanker task scores between Chemo and Control groups.

### 3.4 Association of Cognitive Control with Intrinsic Brain Networks

To differentiate task-evoked responses between the Chemo and Control groups, we extracted activation and deactivation patterns for congruent, incongruent, and incongruent > congruent contrasts at the subject level within task-positive (DAN) and task-negative (DMN) networks. This was done by computing the mean across all voxels that met the criteria: greater than or less than the t-statistic threshold of 1 in the DAN and DMN, respectively. The threshold was chosen as a compromise to avoid including all activated/deactivated voxels (|t|>1) and to avoid being too stringent (|t|>2), which would leave too few voxels per subject within the DAN/DMN mask. Although the results did not reach statistical significance, we observed greater activation in the DAN and greater deactivation in the DMN across all contrasts in the Control group than in the Chemo group (**Figure S1**).

## 4 Discussion

### Reaction time

The chemotherapy group showed significantly slower reaction time on both congruent and incongruent trials compared to the control group, but no differences in accuracy, indicating that survivors require additional time to achieve comparable performance levels. These findings align with prior studies (Nordhjem et al. 2026; Chendan Zhou et al. 2020b; Jacola et al. 2016; Stefancin et al. 2020) that similarly demonstrated that pediatric cancer survivors treated with chemotherapy exhibited longer response times. Together, these studies suggest inefficient cognitive processing rather than impaired response control.

### Hypoactivation in Cognitive Control Networks

The hypoactivation in the bilateral inferior frontal gyri (IFG), the left middle temporal gyrus (MTG), and the right insula during incongruent trials in the chemotherapy group is noteworthy. These regions are core components of the cognitive control network, with the IFG playing a critical role in response inhibition and conflict resolution during flanker tasks (Tops and Boksem 2011; Cai et al. 2014). Neuroimaging studies demonstrate that gray matter volume in bilateral prefrontal gyri, left insula, and inferior temporal gyrus correlates positively with interference inhibition efficiency on flanker tasks (Chen et al. 2015). The insula, as part of the salience network, is essential for detecting behaviorally relevant stimuli and switching between task-positive and task-negative networks; the right anterior insula shows particular importance for detecting behaviorally salient events, while the right IFG is more involved in implementing inhibitory control (Eggert et al. 2025; Kim et al. 2025; Tops and Boksem 2011; Cai et al. 2014). Reduced activation in these regions may reflect diminished neural efficiency in conflict detection and resolution, consistent with meta-analytic findings showing that ALL survivors develop functional changes, particularly in frontal regions (Chendan Zhou et al. 2020b).

### Network-Level Alterations and Striatal-Frontal Dysfunction

The incongruent > congruent contrast revealed that controls showed greater activation in the bilateral caudate, right superior temporal gyrus/middle temporal gyrus, and right postcentral gyrus. In contrast, the chemotherapy group showed greater activation in the bilateral cerebellum and left inferior temporal gyrus. The caudate is integral to cognitive flexibility and action selection (Grahn, Parkinson, and Owen 2008); white-matter microstructure in and around the basal ganglia predicts attention-switching performance, with local white matter projecting to regions of the prefrontal cortex and thalamus via fronto-striato-thalamic circuits (Van Schouwenburg et al. 2014). Furthermore, caudate volume predicts modulation of conflict monitoring during tasks that require parallel conflict processing and flexible action adaptation (Grahn, Parkinson, and Owen 2008). The reduced caudate engagement during conflict processing suggests impaired striatal-frontal circuit function in chemotherapy-exposed patients, consistent with evidence that fronto-striatal circuits are critical for cognitive flexibility (Uddin 2021; Van Schouwenburg et al. 2014).

The attenuated activation in the dorsal attention network (DAN) and reduced deactivation in the default mode network (DMN) across all task contrasts represent a fundamental disruption in network dynamics. The DAN, encompassing the dorsolateral prefrontal cortex and posterior parietal regions, supports externally-directed attention and is typically activated during demanding cognitive tasks (Visintin et al. 2015). The DMN, conversely, is normally suppressed during goal-directed cognition (Anderson, Folk, and Courtney 2016). The failure to adequately activate the DAN and deactivate the DMN suggests impaired network segregation and reduced capacity for cognitive control. This pattern is consistent with computational models demonstrating that activation of the frontoparietal control network and the DAN increases global synchrony and decreases metastability, whereas activation of the DMN has the opposite effects, suggesting that the balance between these networks controls global brain metastability and attentional focus (Hellyer et al. 2014). The chemotherapy group’s inability to achieve this balance may explain their reduced cognitive flexibility and the observed behavioral deficits on standardized testing.

### Compensatory Mechanisms and Alternative Neural Pathways

The associations between the incongruent > congruent contrast and reaction time revealed greater activation in right orbitofrontal cortex (OFC), bilateral inferior temporal gyrus, and right fusiform gyrus in the chemotherapy group, alongside greater deactivations in regions including the left temporal pole. The OFC is involved in value-based decision-making and behavioral flexibility, and its increased recruitment may represent a compensatory mechanism to maintain task performance despite underlying neural inefficiency. The chemotherapy group showed hyperactivation in the fusiform gyrus and temporal regions, which are typically associated with visual processing and semantic encoding. This pattern mirrors recent neuroimaging findings showing that chemotherapy induces microvascular and microstructural brain abnormalities in regions involved in cognition and sensory processing, with decreased perfusion fraction and increased free water fraction associated with poor performance on cognitive tests assessing inhibition and processing speed (McKenna et al. 2026). The concurrent hypoactivation in prefrontal control regions suggests a shift from top-down executive control to bottom-up perceptual processing strategies.

The bilateral cerebellar hyperactivation during the incongruent > congruent contrast in the chemotherapy group is particularly intriguing. The cerebellum contributes to cognitive control through its connections with the prefrontal cortex, and its increased recruitment may represent an alternative neural pathway for conflict resolution when frontoparietal circuits are compromised. This interpretation is supported by evidence that cerebellar activation correlates with cognitive performance in various neurological conditions characterized by frontal dysfunction, and aligns with the compensatory activation theory proposed by previous studies (Robinson et al. 2010; Fellah et al. 2019). wherein survivors recruit additional neural resources to achieve comparable behavioral outcomes.

### Limitations and Future Directions

Several limitations warrant consideration. The cross-sectional design and the absence of pre-treatment baseline neuroimaging limits our ability to distinguish treatment-induced changes from confounders such as pre-existing vulnerabilities. The relatively modest sample size may have limited statistical power to detect subtle group differences, particularly in network-level analyses comparing DAN activation and DMN deactivation patterns, which showed consistent trends but did not reach statistical significance. Heterogeneity in treatment protocols, including variations in chemotherapy agents and intrathecal methotrexate exposure, may contribute to variability in outcomes. Additional fMRI tasks beyond the flanker task would further characterize other domain-specific neural alterations.

## 5 Conclusions

Pediatric cancer survivors treated with chemotherapy demonstrate slower reaction times, hypoactivation in cognitive control networks, impaired striatal-frontal circuit function, and disrupted network dynamics. Compensatory mechanisms, including cerebellar and orbitofrontal recruitment, partially maintain performance. These findings highlight neuroplasticity-based interventions as promising approaches to ameliorate chemotherapy-related cognitive impairments in this vulnerable population.

## Author Contributions

SN contributed to data conceptualization, analysis, interpretation, initial manuscript drafting, and guidance of the manuscript review process. SN and SH contributed to data acquisition. SN, SH, and FM contributed to the interpretation of the data and critically reviewed the manuscript for important intellectual content. TD conceptualized and designed the study, guided data analyses and interpretation, and critically reviewed the manuscript.

## DATA AVAILABILITY STATEMENT

Data are available for sharing upon request.

## Abbreviations

fMRI: Functional magnetic resonance imaging
ALL: Acute lymphoblastic leukemia
CRCI: Chemotherapy-related cognitive impairment
DAN: Dorsal attention network
DMN: Default mode network
MTG: Middle temporal gyrus
ITG: Inferior temporal gyrus
STG: Superior temporal gyrus
SMA: Supplementary motor area
IFG: Inferior frontal gyrus
OFC: Orbitofrontal cortex
SFG: Superior frontal gyrus

## References

Anderson, Brian A, Charles L Folk, and Susan M Courtney. 2016. “Neural mechanisms of goal-contingent task disengagement: Response-irrelevant stimuli activate the default mode network.” Cortex 81: 221–230.

Cai, Weidong, Srikanth Ryali, Tianwen Chen, Chiang-Shan R Li, and Vinod Menon. 2014. “Dissociable roles of right inferior frontal cortex and anterior insula in inhibitory control: evidence from intrinsic and task-related functional parcellation, connectivity, and response profile analyses across multiple datasets.” Journal of Neuroscience 34 (44): 14652–14667.

Chen, Changming, Jiemin Yang, Jiayu Lai, Hong Li, Jiajin Yuan, and Najam ul Hasan Abbasi. 2015. “Correlating gray matter volume with individual difference in the flanker interference effect.” PloS one 10 (8): e0136877.

Cheung, Yin Ting, and Kevin R Krull. 2015. “Neurocognitive outcomes in long-term survivors of childhood acute lymphoblastic leukemia treated on contemporary treatment protocols: A systematic review.” Neuroscience & Biobehavioral Reviews 53: 108–120.

Chughtai, Shahzaib, David Doyle, Swathi Tata, Dhiya Ram, and Irfan Oymagil. 2025. “Chemotherapy-induced cognitive impairment: Mechanisms, emerging biomarkers, and therapeutic interventions.” Biochemical and Biophysical Research Communications: 152456.

Conklin, Heather M, Robert J Ogg, Jason M Ashford, Matthew A Scoggins, Ping Zou, Kellie N Clark, Karen Martin-Elbahesh, Kristina K Hardy, Thomas E Merchant, and Sima Jeha. 2015. “Computerized cognitive training for amelioration of cognitive late effects among childhood cancer survivors: a randomized controlled trial.” Journal of clinical oncology 33 (33): 3894–3902.

Edelmann, M. N., R. J. Ogg, M. A. Scoggins, T. M. Brinkman, N. D. Sabin, C. H. Pui, D. K. Srivastava, L. L. Robison, M. M. Hudson, and K. R. Krull. 2013. “Dexamethasone exposure and memory function in adult survivors of childhood acute lymphoblastic leukemia: A report from the SJLIFE cohort.” Pediatr Blood Cancer 60 (11): 1778–84. 10.1002/pbc.24644. https://www.ncbi.nlm.nih.gov/pubmed/23775832.

Eggert, Patrick, Moritz Mückschel, Nasibeh Talebi, Christian Beste, and Filippo Ghin. 2025. “On the role of the insula cortex in inhibitory control: insights from alpha and theta directed connectivity dynamics.” Cerebral Cortex 35 (10): bhaf292.

Fellah, S., Y. T. Cheung, M. A. Scoggins, P. Zou, N. D. Sabin, C. H. Pui, L. L. Robison, M. M. Hudson, R. J. Ogg, and K. R. Krull. 2019. “Brain Activity Associated With Attention Deficits Following Chemotherapy for Childhood Acute Lymphoblastic Leukemia.” J Natl Cancer Inst 111 (2): 201–209. 10.1093/jnci/djy089. https://pmc.ncbi.nlm.nih.gov/articles/PMC6376909/.

Gajjar, Amar, Anita Mahajan, Tejus Bale, Daniel C Bowers, Liz Canan, Susan Chi, Andrew Cluster, Kenneth Cohen, Bonnie Cole, and Scott Coven. 2025. “Pediatric central nervous system cancers, version 2.2025, NCCN clinical practice guidelines in oncology.” Journal of the National Comprehensive Cancer Network 23 (3): 113–130.

Gandy, K., Y. Sapkota, M. A. Scoggins, L. M. Jacola, T. R. Koscik, M. M. Hudson, C. H. Pui, K. R. Krull, and E. van der Plas. 2023. “Genetic variants, neurocognitive outcomes, and functional neuroimaging in survivors of childhood acute lymphoblastic leukemia.” JNCI Cancer Spectr 7 (4). 10.1093/jncics/pkad039. https://www.ncbi.nlm.nih.gov/pubmed/37285328.

Gandy, K., M. A. Scoggins, L. M. Jacola, M. Litten, W. E. Reddick, and K. R. Krull. 2021. “Structural and Functional Brain Imaging in Long-Term Survivors of Childhood Acute Lymphoblastic Leukemia Treated With Chemotherapy: A Systematic Review.” JNCI Cancer Spectr 5 (5). 10.1093/jncics/pkab069. https://www.ncbi.nlm.nih.gov/pubmed/34514328.

Gandy, K., M. A. Scoggins, N. Phillips, E. van der Plas, S. Fellah, L. M. Jacola, C. H. Pui, M. M. Hudson, W. E. Reddick, R. Sitaram, and K. R. Krull. 2022. “Sex-Based Differences in Functional Brain Activity During Working Memory in Survivors of Pediatric Acute Lymphoblastic Leukemia.” JNCI Cancer Spectr 6 (2). 10.1093/jncics/pkac026. https://www.ncbi.nlm.nih.gov/pubmed/35603857.

Grahn, Jessica A, John A Parkinson, and Adrian M Owen. 2008. “The cognitive functions of the caudate nucleus.” Progress in neurobiology 86 (3): 141–155.

Hellyer, Peter J, Murray Shanahan, Gregory Scott, Richard JS Wise, David J Sharp, and Robert Leech. 2014. “The control of global brain dynamics: opposing actions of frontoparietal control and default mode networks on attention.” Journal of Neuroscience 34 (2): 451–461.

Hooke, M. C., M. A. Mathiason, A. S. Kunin-Batson, A. Blommer, J. Hutter, P. A. Mitby, I. M. Moore, S. Whitman, O. Taylor, M. E. Scheurer, and M. J. Hockenberry. 2021. “Biomarkers and Cognitive Function in Children and Adolescents During Maintenance Therapy for Leukemia.” Oncol Nurs Forum 48 (6): 623–633. 10.1188/21.ONF.623-633. https://www.ncbi.nlm.nih.gov/pubmed/34673759.

Hunger, Stephen P, and Charles G Mullighan. 2015. “Acute lymphoblastic leukemia in children.” New England Journal of Medicine 373 (16): 1541–1552.

Inaba, Hiroto, David Teachey, Colleen Annesley, Sandeep Batra, Jill Beck, Susan Colace, Stacy Cooper, Mari Dallas, Satiro De Oliveira, and Kara Kelly. 2025. “Pediatric acute lymphoblastic leukemia, version 2.2025, NCCN clinical practice guidelines in oncology.” Journal of the National Comprehensive Cancer Network 23 (2): 41–62.

Jacola, Lisa M, Kevin R Krull, Ching-Hon Pui, Deqing Pei, Cheng Cheng, Wilburn E Reddick, and Heather M Conklin. 2016. “Longitudinal assessment of neurocognitive outcomes in survivors of childhood acute lymphoblastic leukemia treated on a contemporary chemotherapy protocol.” Journal of Clinical Oncology 34 (11): 1239–1247.

Jacola, Lisa M, Rachel K Peterson, Kaitlin A Oswald-McCloskey, Angela Sekely, Donald J Mabbott, and Kim Edelstein. 2025. “Neuropsychological function in childhood cancer patients and adult survivors of childhood cancer.” Journal of Clinical and Experimental Neuropsychology 47 (8): 788–805.

Jenkinson, Mark, Christian F Beckmann, Timothy EJ Behrens, Mark W Woolrich, and Stephen M Smith. 2012. “Fsl.” Neuroimage 62 (2): 782–790.

Kim, Jiyea, Narae Kim, Yeeun Kim, Bumhee Park, and Min Hyeon Park. 2025. “The association between fronto-insular interaction and executive function during childhood and adolescence in a resting state fMRI study.” Scientific Reports 15 (1): 30096.

Malard, Florent, and Mohamad Mohty. 2020. “Acute lymphoblastic leukaemia.” The Lancet 395 (10230): 1146–1162.

McDermott, Timothy J, Alex I Wiesman, Amy L Proskovec, Elizabeth Heinrichs-Graham, and Tony W Wilson. 2017. “Spatiotemporal oscillatory dynamics of visual selective attention during a flanker task.” Neuroimage 156: 277–285.

McKenna, F., S. Nandi, S. S. Henry, S. Nayak, R. Fleysher, and T. Q. Duong. 2026. “Microvascular and microstructural brain abnormalities in paediatric haematological cancer survivors are related to cognitive deficits: An IVIM-FWI MRI study.” Br J Haematol 208 (2): 608–619. 10.1111/bjh.70300. https://www.ncbi.nlm.nih.gov/pubmed/41508157.

Monje, Michelle, Moriah E Thomason, Laura Rigolo, Yalin Wang, Deborah P Waber, Stephen E Sallan, and Alexandra J Golby. 2013. “Functional and structural differences in the hippocampus associated with memory deficits in adult survivors of acute lymphoblastic leukemia.” Pediatric blood & cancer 60 (2): 293–300.

Nordhjem, Barbara Johanne Thomas, Liv Andrés-Jensen, Kristian Mielke Christensen, Marianne Helenius, Birthe Lykke Thomsen, Ingrid Tonning Olsson, Hanne Bækgaard Larsen, and Lisa Lyngsie Hjalgrim. 2026. “Neurocognitive outcomes in survivors of ALL: Risk patterns and individual profiles in a single-protocol cohort.” Journal of the International Neuropsychological Society: 1–12.

Park, Yongkyu, Nirajan K. C, Jeremy Willekens, Chadni Patel, Beth A Savage, Haiqun Lin, Alysta Paneque, Robert Daly, Alexandra Thrope, and Melissa A Burns. 2025. “Treatment-related changes in cerebrospinal fluid markers of oxidative stress and neurodegeneration during therapy for childhood acute lymphoblastic leukemia.” Cancer Epidemiology, Biomarkers & Prevention 34 (11): 2015–2024.

Robinson, K. E., K. L. Livesay, L. K. Campbell, M. Scaduto, C. J. Cannistraci, A. W. Anderson, J. A. Whitlock, and B. E. Compas. 2010. “Working memory in survivors of childhood acute lymphocytic leukemia: functional neuroimaging analyses.” Pediatr Blood Cancer 54 (4): 585–90. 10.1002/pbc.22362. https://pmc.ncbi.nlm.nih.gov/articles/PMC2901833/.

Rolls, Edmund T, Chu-Chung Huang, Ching-Po Lin, Jianfeng Feng, and Marc Joliot. 2020. “Automated anatomical labelling atlas 3.” Neuroimage 206: 116189.

Shono, Yusuke, Berivan Ece, Emily H Ho, Aaron J Kaat, Erica M LaForte, Ezgi Ayturk, and Richard Gershon. 2024. “A comparison of scoring algorithms for the NIH Toolbox executive function tasks in a US norming sample.” Psychological assessment 36 (12): 760.

Smith, Stephen M, Mark Jenkinson, Mark W Woolrich, Christian F Beckmann, Timothy EJ Behrens, Heidi Johansen-Berg, Peter R Bannister, Marilena De Luca, Ivana Drobnjak, and David E Flitney. 2004. “Advances in functional and structural MR image analysis and implementation as FSL.” Neuroimage 23: S208–S219.

Stefancin, P., C. Cahaney, R. I. Parker, T. Preston, K. Coulehan, L. Hogan, and T. Q. Duong. 2020. “Neural correlates of working memory function in pediatric cancer survivors treated with chemotherapy: an fMRI study.” NMR Biomed 33 (6): e4296. 10.1002/nbm.4296. https://www.ncbi.nlm.nih.gov/pubmed/32215994.

Tops, Mattie, and Maarten AS Boksem. 2011. “A potential role of the inferior frontal gyrus and anterior insula in cognitive control, brain rhythms, and event-related potentials.” Frontiers in psychology 2: 330.

Uddin, Lucina Q. 2021. “Cognitive and behavioural flexibility: neural mechanisms and clinical considerations.” Nature reviews neuroscience 22 (3): 167–179.

Van Schouwenburg, Martine Rinske, A Marten H Onnink, Niels Ter Huurne, Cees C Kan, Marcel P Zwiers, Martine Hoogman, Barbara Franke, Jan K Buitelaar, and Roshan Cools. 2014. “Cognitive flexibility depends on white matter microstructure of the basal ganglia.” Neuropsychologia 53: 171–177.

Visintin, Eleonora, Chiara De Panfilis, Camilla Antonucci, Cinzia Capecci, Carlo Marchesi, and Fabio Sambataro. 2015. “Parsing the intrinsic networks underlying attention: a resting state study.” Behavioural brain research 278: 315–322.

Weintraub, Sandra, Sureyya S Dikmen, Robert K Heaton, David S Tulsky, Philip D Zelazo, Patricia J Bauer, Noelle E Carlozzi, Jerry Slotkin, David Blitz, and Kathleen Wallner-Allen. 2013. “Cognition assessment using the NIH Toolbox.” Neurology 80 (11_supplement_3): S54-S64.

Zhou, C., Y. Zhuang, X. Lin, A. D. Michelson, and A. Zhang. 2020a. “Changes in neurocognitive function and central nervous system structure in childhood acute lymphoblastic leukaemia survivors after treatment: a meta-analysis.” Br J Haematol 188 (6): 945–961. 10.1111/bjh.16279. https://www.ncbi.nlm.nih.gov/pubmed/31823355.

Zhou, Chendan, Yong Zhuang, Xingjie Lin, Alan D Michelson, and Aijun Zhang. 2020b. “Changes in neurocognitive function and central nervous system structure in childhood acute lymphoblastic leukaemia survivors after treatment: a meta analysis.” British Journal of Haematology 188 (6): 945–961.

Zhu, David C, Rose T Zacks, and Jill M Slade. 2010. “Brain activation during interference resolution in young and older adults: an fMRI study.” Neuroimage 50 (2): 810–817.

