## Supplemental Material for "Cognitive Control in Pediatric Cancer Survivors: A task-fMRI study"

**Supplementary Materials for** “**Cognitive control in Pediatric Cancer Survivors: A task-fMRI study.”**

**Table S1: Brain regions showing significant activation/deactivation for the (A) Congruent, (B) Incongruent, and (C) Incongruent > Congruent contrasts between Chemo and Control groups**

| **(A)** | **Volume, mm^3^** | **Peak T** | **XYZ** | **Structure** |
| --- | --- | --- | --- | --- |
| Chemo | 5299 | 6.1 | -4.2×18.3×46.2 | Left Supplementary Motor Area |
|  | 38791 | 5.2 | 59.9×-43.7×-6.1 | Right Middle Temporal Gyrus |
|  | 2227 | 5.1 | 62.1×19.8×17.5 | Right Inferior Frontal Operculum |
|  | 33565 | 4.9 | -35.9×-84.2×3.5 | Left Middle Occipital Gyrus |
|  | 1459 | 4.7 | 44.4×-35.5×-21.6 | Right Fusiform Gyrus |
|  | 998 | 4.5 | 17.9×52.2×-17.9 | Right Orbitofrontal Cortex |
|  | 11122 | 4.4 | 45.9×3.5×48.4 | Right Precentral Gyrus |
|  | 266 | -4.1 | -33.7×-16.4×55.8 | Left Precentral Gyrus |
|  | 972 | -4.2 | -13.8×58.1×-1.7 | Left Superior Medial Frontal |
|  | 587 | -4.3 | -44.1×-7.5×-29.7 | Left Inferior Temporal Gyrus |
|  | 209 | -4.4 | 3.9×36.0×-23.8 | Right Rectus |
|  | 2201 | -4.6 | -61.8×-17.8×-0.2 | Left Middle Temporal Gyrus |
| Control | 13922 | 5.3 | -62.5×-29.6×-9.8 | Left Middle Temporal Gyrus |
|  | 12040 | 4.7 | 31.9×55.9×27.8 | Right Middle Frontal Gyrus |
|  | 9788 | 4.3 | -6.4×10.2×52.1 | Left Supplementary Motor Area |
|  | 15697 | 4.3 | -4.2×-70.2×-12.0 | Left Cerebellum |
|  | 2631 | 4.3 | -12.3×36.0×54.3 | Left Superior Frontal Gyrus |
|  | 4295 | 4.2 | -23.4×-6.0×0.5 | Left Pallidum |
|  | 2519 | 4.1 | -27.8×-68.0×61.7 | Left Superior Parietal |
|  | 195 | -3.1 | 24.5×27.9×-12.0 | Right Orbitofrontal Cortex |
|  | 126 | -3.1 | 7.6×-84.2×10.1 | Right Calcarine Cortex |
|  | 161 | -3.5 | 26.0×-0.1×-35.6 | Right ParaHippocampal Gyrus |
|  | 2323 | -4.4 | -18.2×32.3×-15.7 | Left Orbitofrontal Cortex |
|  | 551 | -4.7 | -52.2×-43.7×7.9 | Left Middle Temporal Gyrus |
| Control > Chemo | 567 | 4.5 | -45.5×-9.7×-28.3 | Left Inferior Temporal Gyrus |
|  | 197 | 4.1 | 45.9×-3.8×-28.3 | Right Middle Temporal Gyrus |
|  | 7224 | 4.1 | -16.0×-40.0×62.5 | Left Supplementary Motor Area |
|  | 461 | 3.9 | -61.8×-16.4×-0.2 | Left Middle Temporal Gyrus |
|  | 532 | 3.9 | -16.0×11.7×49.9 | Left Superior Frontal Gyrus |
|  | 1762 | 3.9 | 31.9×53.7×31.5 | Right Middle Frontal Gyrus |
|  | 753 | 3.6 | -2.0×8.0×33.7 | Left Middle Cingulate Cortex |
|  | 182 | -3.1 | 48.1×-64.3×21.9 | Right Middle Temporal Gyrus |
|  | 325 | -3.3 | -3.5×18.3×46.2 | Left Supplementary Motor Area |
|  | 451 | -3.6 | 30.4×30.1×-15.7 | Right Orbitofrontal Cortex |
|  | 1152 | -3.7 | 59.9×-43.7×-5.4 | Right Middle Temporal Gyrus |
|  | 480 | -3.7 | 59.9×24.2×12.3 | Right Inferior Frontal Triangular |

| **(B)** | **Volume, mm^3^** | **Peak T** | **XYZ** | **Structure** |
| --- | --- | --- | --- | --- |
| Chemo | 30512 | 6.1 | -35.9×-84.2×2.0 | Left Middle Occipital Gyrus |
|  | 47623 | 6 | 42.2×-68.0×-6.1 | Right Inferior Temporal Gyrus |
|  | 2793 | 5.3 | -3.5×19.8×46.2 | Left Supplementary Motor Area |
|  | 2021 | 4.4 | 58.5×-41.4×-12.0 | Right Inferior Temporal Gyrus |
|  | 3170 | 4.3 | 26.0×45.6×-16.5 | Right Orbitofrontal Cortex |
|  | 1916 | 4.1 | 26.0×-34.1×7.9 | Right Hippocampus |
|  | 13267 | 4 | 37.8×5.8×58.0 | Right Middle Frontal Gyrus |
|  | 8149 | -4.1 | -16.0×-40.0×63.9 | Right Supplementary Motor Area |
|  | 2865 | -4.2 | -53.6×25.7×10.1 | Left Inferior Frontal Triangular |
|  | 2443 | -4.4 | -16.0×-64.3×-5.4 | Left Lingual Gyrus |
|  | 9964 | -4.7 | -6.4×55.9×20.4 | Left Superior Medial Frontal |
|  | 35577 | -5.3 | -45.5×-34.1×25.6 | Left SupraMarginal Gyrus |
| Control | 184798 | 5.3 | 37.8×-84.2×-0.2 | Right Middle Occipital Gyrus |
|  | 6849 | 4.5 | 26.0×-1.6×16.0 | Right Thalamus |
|  | 1885 | 4 | -30.0×44.1×18.2 | Left Middle Frontal Gyrus |
|  | 5854 | 4 | 29.7×44.1×16.0 | Right Middle Frontal Gyrus |
|  | 670 | 3.8 | 37.8×36.0×5.7 | Right Inferior Frontal Triangular |
|  | 1476 | 3.6 | 48.1×-22.3×16.0 | Right Rolandic Operculum |
|  | 431 | 3.5 | -21.9×52.2×-14.2 | Left Orbitofrontal Cortex |
|  | 1011 | -3.2 | -12.3×47.8×-12.0 | Left Medial OrbitoFrontal Cortex |
|  | 192 | -3.3 | 12.0×47.8×-6.1 | Right Medial OrbitoFrontal Cortex |
|  | 430 | -3.4 | -20.5×32.3×-15.7 | Left Orbitofrontal Cortex |
|  | 2916 | -3.4 | -16.0×-62.1×20.4 | Left Cuneus |
|  | 189 | -3.6 | -50.0×-65.8×24.1 | Left Angular Gyrus |
| Control > Chemo | 39285 | 5.3 | -31.5×-17.8×12.3 | Left Postcentral Gyrus |
|  | 6795 | 4.1 | 38.5×3.5×16.0 | Right Insula |
|  | 818 | 3.9 | 40.0×33.8×5.7 | Right Inferior Frontal Triangular |
|  | 2013 | 3.8 | -5.7×55.9×19.7 | Left Superior Medial Frontal |
|  | 1529 | 3.7 | -64.0×-23.7×-12.0 | Left Middle Temporal Gyrus |
|  | 5224 | 3.7 | -58.1×13.9×2.0 | Left Inferior Frontal Operculum |
|  | 491 | 3.7 | 43.7×-29.6×63.9 | Right Postcentral Gyrus |
|  | 384 | -2.7 | 26.0×-70.2×30.0 | Right Superior Occipital Gyrus |
|  | 91 | -2.8 | -34.5×-16.4×-14.2 | Left Hippocampus |
|  | 651 | -2.9 | 47.4×22.0×32.2 | Right Inferior Frontal Operculum |
|  | 461 | -3 | 62.1×-44.4×-16.5 | Right Inferior Temporal Gyrus |
|  | 274 | -3.5 | 48.1×-63.6×21.9 | Right Middle Temporal Gyrus |

| **(C)** | **Volume, mm^3^** | **Peak T** | **XYZ** | **Structure** |
| --- | --- | --- | --- | --- |
| Chemo | 465 | 3.9 | 28.2×-51.8×38.1 | Right Inferior Parietal Gyrus |
|  | 2670 | 3.9 | -3.5×-34.1×-35.6 | Left Cerebellum |
|  | 286 | 3.8 | 20.1×11.7×38.1 | Right Superior Frontal Gyrus |
|  | 272 | 3.4 | 40.0×-73.9×7.9 | Right Middle Occipital Gyrus |
|  | 85 | 3.4 | -12.3×2.1×-20.1 | Left ParaHippocampal |
|  | 504 | 3.3 | -6.4×-42.2×-26.0 | Left Cerebellum |
|  | 105 | 3.2 | 3.9×-96.0×2.0 | Left Calcarine |
|  | 6314 | -4.2 | -18.2×2.1×12.3 | Left Putamen |
|  | 565 | -4.2 | 8.3×19.8×-8.3 | Right Olfactory |
|  | 827 | -4.3 | -27.8×-11.9×-0.2 | Left Putamen |
|  | 6161 | -4.3 | -30.0×45.6×39.6 | Left Superior Frontal Gyrus |
|  | 3575 | -6.1 | -11.6×22.0×-9.8 | Left Medial OrbitoFrontal Cortex |
| Control | 9037 | 4.5 | 23.8×-60.6×49.9 | Right Superior Parietal |
|  | 4842 | 4 | -24.1×4.3×54.3 | Left Superior Frontal Gyrus |
|  | 3227 | 3.9 | 49.6×-77.6×10.1 | Right Middle Temporal Gyrus |
|  | 691 | 3.9 | -17.5×8.0×10.1 | Left Putamen |
|  | 2067 | 3.7 | 28.2×-3.8×51.4 | Right Superior Frontal Gyrus |
|  | 785 | 3.7 | -31.5×-9.7×0.5 | Left Putamen |
|  | 547 | 3.7 | -30.0×-82.0×17.5 | Left Middle Occipital Gyrus |
|  | 838 | -3.3 | 54.0×-18.6×-15.7 | Right Middle Temporal Gyrus |
|  | 484 | -3.4 | -46.3×-6.0×-21.6 | Left Middle Temporal Gyrus |
|  | 862 | -3.7 | 17.9×55.9×35.9 | Right Superior Frontal Gyrus |
|  | 769 | -3.9 | 22.3×64.0×9.4 | Right Superior Frontal Gyrus |
|  | 375 | -4 | -38.2×-76.1×-23.8 | Left Cerebellum |
| Control > Chemo | 6609 | 5.1 | -27.8×-11.9×-0.2 | Left Putamen |
|  | 1297 | 4.7 | -12.3×22.0×-8.3 | Left Caudate |
|  | 1949 | 4.2 | 26.0×-11.9×16.0 | Right Caudate |
|  | 845 | 3.9 | 57.7×-54.0×4.2 | Right Middle Temporal Gyrus |
|  | 282 | 3.7 | 51.8×-11.9×0.5 | Right Superior Temporal Gyrus |
|  | 220 | 3.4 | 45.9×-28.2×63.9 | Right Postcentral Gyrus |
|  | 403 | 3.3 | 55.5×15.3×-12.0 | Right Temporal Pole |
|  | 132 | -2.7 | 31.9×-8.3×-16.5 | Right Hippocampus |
|  | 269 | -2.8 | 20.1×67.7×7.9 | Right Superior Frontal Gyrus |
|  | 171 | -2.9 | -53.6×-57.7×-29.7 | Left Inferior Temporal Gyrus |
|  | 327 | -3.5 | 20.1×-87.9×-21.6 | Right Cerebellum |
|  | 604 | -3.8 | -35.9×-78.3×-23.8 | Left Cerebellum |

**Table S2: Brain regions showing significant association of Incongruent > Congruent contrast with (A) reaction time and (B) NIH flanker task scores between Chemo and Control groups.**

| **(A)** | **Volume, mm^3^** | **Peak T** | **XYZ** | **Structure** |
| --- | --- | --- | --- | --- |
| Chemo | 1273 | 5.4 | 10.5×41.9×-27.5 | Right Medial OrbitoFrontal Cortex |
|  | 403 | 5.2 | -24.1×-43.7×-35.6 | Left Cerebellum |
|  | 3872 | 5 | -44.1×-5.3×-46.0 | Left Fusiform Gyrus |
|  | 2796 | 4.6 | 22.3×44.1×16.0 | Right Anterior Cingulate Cortex |
|  | 2094 | 4.4 | 38.5×-2.4×-46.0 | Right Inferior Temporal Gyrus |
|  | 6897 | 4.4 | -24.1×-31.9×21.9 | Left Anterior Cingulate Cortex |
|  | 1417 | 4.3 | 36.3×-42.2×27.8 | Right SupraMarginal Gyrus |
|  | 531 | -4.5 | 56.2×-3.8×-9.8 | Right Superior Temporal Gyrus |
|  | 3870 | -4.5 | -26.4×51.5×-12.0 | Left Middle Frontal Gyrus |
|  | 710 | -4.7 | 54.0×-56.2×-34.2 | Right Cerebellum |
|  | 7625 | -6.1 | 45.9×-1.6×-26.0 | Right Middle Temporal Gyrus |
|  | 1124 | -6.2 | 9.8×72.1×5.7 | Right Superior Medial Frontal |
| Control | 172 | 5.1 | -30.0×-34.1×-34.2 | Left Cerebellum |
|  | 1348 | 4.2 | 33.4×-59.9×49.9 | Right Angular Gyrus |
|  | 1497 | 4.1 | 8.3×-57.7×-20.1 | Right Cerebellum |
|  | 727 | 4 | 28.2×-72.4×30.0 | Right Middle Occipital Gyrus |
|  | 677 | 3.9 | -41.8×-9.7×34.4 | Left Postcentral Gyrus |
|  | 93 | 3.9 | 63.6×-23.7×-28.3 | Right Inferior Temporal Gyrus |
|  | 2424 | 3.9 | 49.6×4.3×26.3 | Right Precentral Gyrus |
|  | 615 | -3.3 | -0.5×-58.4×30.0 | Left Precuneus |
|  | 414 | -3.4 | 6.1×-34.1×35.9 | Right Middle Cingulate |
|  | 141 | -3.6 | 26.0×-79.8×-26.0 | Right Crus Cerebellum |
|  | 225 | -3.8 | -13.8×16.1×-12.0 | Left Rectus |
|  | 1026 | -3.8 | -4.2×64.0×12.3 | Left Superior Medial Frontal |
| Control > Chemo | 904 | 5.3 | 9.8×72.1×5.7 | Right Superior Frontal Gyrus |
|  | 4340 | 5 | 59.9×8.0×-22.4 | Right Middle Temporal Pole |
|  | 1730 | 4.8 | 21.6×-8.3×-16.5 | Right Hippocampus |
|  | 1893 | 4.5 | -18.2×-11.9×-27.5 | Left ParaHippocampal Gyrus |
|  | 479 | 4.1 | 14.2×-17.8×-0.2 | Right Thalamus |
|  | 907 | 4.1 | 12.0×-54.0×-20.1 | Right Cerebellum |
|  | 677 | 3.8 | -41.8×-9.7×34.4 | Left Postcentral |
|  | 172 | -3.3 | 8.3×-88.6×-34.2 | Right Crus Cerebellum |
|  | 100 | -3.3 | -44.1×11.7×-12.0 | Left Insula |
|  | 306 | -3.4 | -56.6×-55.5×-20.1 | Left Inferior Temporal |
|  | 269 | -3.7 | 57.7×-37.8×-27.5 | Right Inferior Temporal |
|  | 870 | -3.8 | 37.8×-1.6×-46.0 | Right Inferior Temporal |

| **(B)** | **Volume, mm^3^** | **Peak T** | **XYZ** | **Structure** |
| --- | --- | --- | --- | --- |
| Chemo | 2183 | 5.5 | 16.4×-6.0×-28.3 | Right Fusiform Gyrus |
|  | 5232 | 5.4 | -61.8×-17.8×-28.3 | Left Inferior Temporal Gyrus |
|  | 4215 | 4.4 | -41.8×4.3×-46.0 | Left Fusiform Gyrus |
|  | 578 | 4.3 | -27.8×11.7×13.8 | Left Putamen |
|  | 2181 | 4.2 | 3.9×10.2×-22.4 | Right Rectus |
|  | 352 | 4.1 | 13.5×45.6×-26.0 | Right Medial OrbitoFrontal Cortex |
|  | 2026 | 3.9 | 49.6×-2.4×-46.0 | Right Inferior Temporal Gyrus |
|  | 769 | -4.8 | -33.7×23.5×-35.6 | Left Temporal Pole |
|  | 2055 | -4.9 | 20.1×41.9×-3.9 | Right Medial OrbitoFrontal Cortex |
|  | 501 | -4.9 | -17.5×67.7×19.7 | Left Superior Frontal Gyrus |
|  | 3067 | -5.3 | 12.0×65.5×27.8 | Right Superior Medial Frontal |
|  | 509 | -5.6 | -13.8×4.3×-3.9 | Left Pallidum |
| Control | 335 | 4.3 | -67.7×-36.3×4.2 | Left Middle Temporal Gyrus |
|  | 574 | 4.2 | -65.4×-11.9×-3.9 | Left Middle Temporal Gyrus |
|  | 275 | 4.1 | -58.1×6.5×-27.5 | Left Middle Temporal Gyrus |
|  | 134 | 3.4 | 45.9×-12.7×33.7 | Right Postcentral Gyrus |
|  | 294 | 3.4 | -41.8×10.2×23.4 | Left Inferior Frontal Triangular |
|  | 1284 | -4.3 | 48.1×-59.9×-19.4 | Right Inferior Temporal Gyrus |
|  | 657 | -4.4 | 31.9×54.4×5.7 | Right Superior Frontal Gyrus |
|  | 661 | -4.7 | 28.2×53.7×24.1 | Right Superior Frontal Gyrus |
|  | 3026 | -4.9 | -7.9×-80.5×-19.4 | Left Crus Cerebellum |
|  | 2860 | -5 | 26.0×-54.0×53.6 | Right Inferior Parietal Lobe |
| Control > Chemo | 528 | 4.5 | 45.9×-11.9×33.7 | Right Postcentral |
|  | 452 | 4.2 | -38.2×22.0×-37.8 | Left Middle Temporal Pole |
|  | 440 | 4 | -65.4×-11.9×-6.1 | Left Middle Temporal |
|  | 230 | 3.9 | -67.7×-35.5×4.2 | Left Middle Temporal |
|  | 291 | 3.5 | -16.0×2.1×-3.9 | Left Pallidum |
|  | 230 | -3.6 | 59.9×8.0×4.2 | Right Rolandic Operculum |
|  | 128 | -3.7 | -55.9×-48.1×-29.7 | Left Cerebellum_Crus |
|  | 311 | -3.9 | 28.2×52.2×24.1 | Right Superior Frontal Gyrus |
|  | 1332 | -4.1 | -32.3×-71.7×-22.4 | Left Cerebellum |
|  | 2264 | -4.5 | 26.0×-54.0×53.6 | Right Inferior Parietal Lobe |

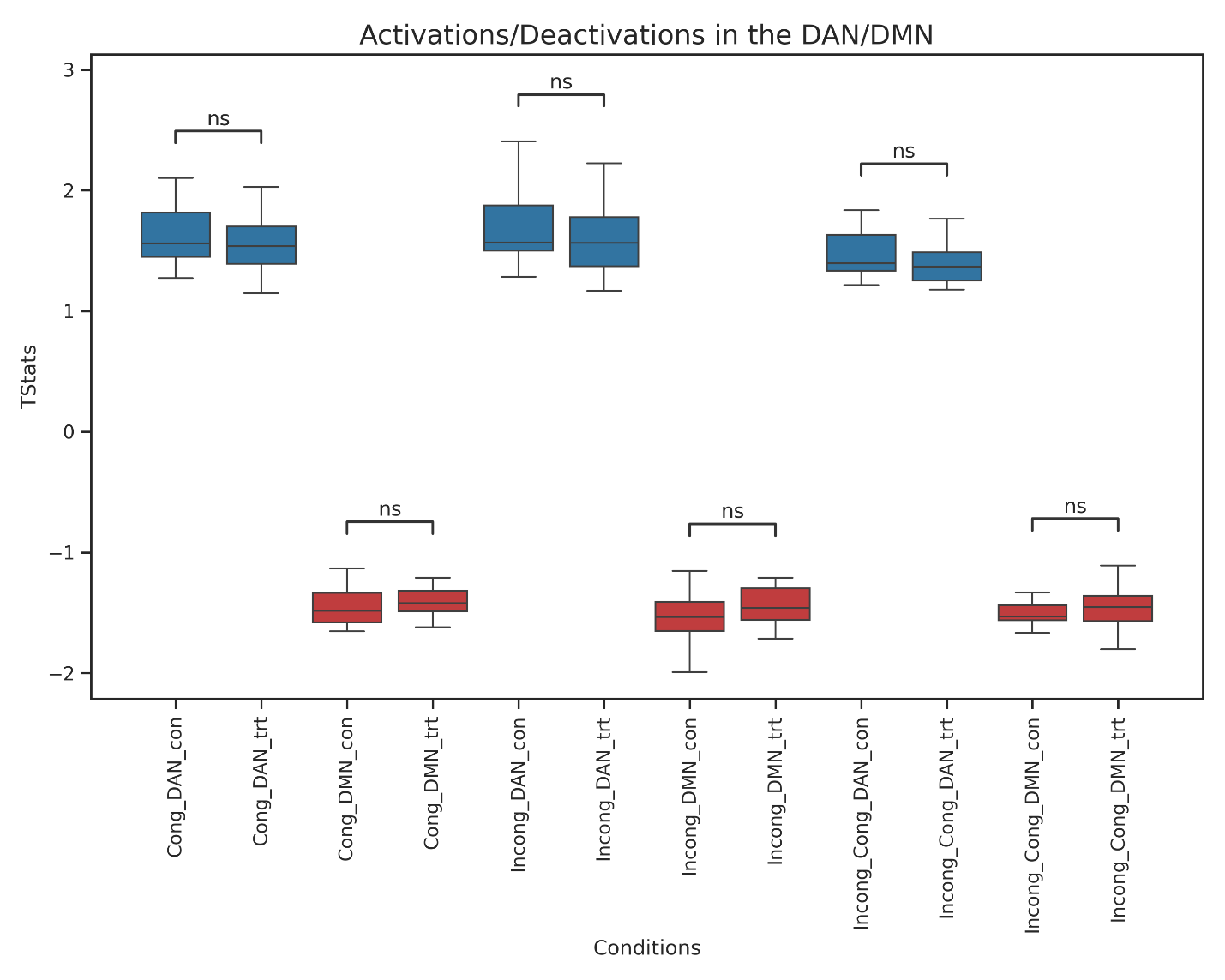

 **Figure S1:** ROI analyses within DAN and DMN networks across Chemo and Control groups. (Top) Boxplots display the mean activation within DAN in blue and (bottom) mean deactivation within DMN in red. Subject-level DAN and DMN values are generated with a t-statistic threshold of t > 1 and t < -1, respectively. Note that controls are abbreviated as ‘con’ and chemo is abbreviated as ‘trt’.
